# Neutralization heterogeneity of United Kingdom and South-African SARS-CoV-2 variants in BNT162b2-vaccinated or convalescent COVID-19 healthcare workers

**DOI:** 10.1101/2021.03.05.434089

**Authors:** Stéphane Marot, Isabelle Malet, Aude Jary, Valentin Leducq, Basma Abdi, Elisa Teyssou, Cathia Soulie, Marc Wirden, Christophe Rodriguez, Slim Fourati, Jean-Michel Pawlotsky, David Boutolleau, Sonia Burrel, Vincent Calvez, Anne-Geneviève Marcelin

**Author notes:** Corresponding author. Stephane Marot, Department of Virology, Pitié Salpêtrière Hospital, AP-HP, CERVI, 83 Boulevard de l’Hôpital, 75013, Paris, France. Alternate corresponding author: Anne-Geneviève Marcelin, Department of Virology, Pitié Salpêtrière Hospital, AP-HP, CERVI, 83 Boulevard de l’Hôpital, 75013, Paris, France.

## Abstract

There are concerns about neutralizing antibodies (NAb) potency against the newly emerged VOC202012/01 (UK) and 501Y.V2 (SA) SARS-CoV-2 variants in mRNA-vaccinated subjects and in recovered COVID-19 patients. We used a viral neutralization test with a strict 100% neutralizing criterion on UK and SA clinical isolates in comparison with a globally distributed D614G SARS-CoV-2 strain. In two doses BNT162b2-vaccinated healthcare workers (HCW), despite heterogeneity in neutralizing capacity against the three SARS-CoV-2 strains, most of the sera harbored at least a NAb titer ≥ 1:10 suggesting a certain humoral protection activity either on UK or SA variants. However, six months after mild forms of COVID-19, an important proportion of HCW displayed no neutralizing activity against SA strain. This result supports strong recommendations for vaccination of previously infected subjects.

## INTRODUCTION

In the gene encoding the Spike (S) protein of SARS-CoV-2, various mutations have been reported[1,2] and recently, the United Kingdom (UK) and South Africa (SA) have faced a rapid increase in COVID-19 cases mediated by the emergence of new variants (VOC-202012/01 for UK and 501Y.V2 for SA)[3,4]. The spreading of these variants has increased rapidly in other countries and recent observations suggests that they are significantly more transmissible than previously circulating variants. It is still not fully known if the pathogenicity is either increased, although some elements have been recently released with likely enhanced disease severity for the UK strain[5].

These variants harbor a specific pattern of deletion and mutations including amino-acid replacements at key sites in the S Receptor Binding Domain (RBD) (K417N, E484K, N501Y for the SA strain and only N501Y for the UK strain). In the era of the COVID-19 vaccination, the question remained whether these variants could escape the neutralizing response elicited by mRNA-vaccine. Two recent studies performed on engineered SARS-CoV-2 viruses containing only some mutations from the newly emerged UK and SA variants showed weaker neutralization capacity of vaccine-elicited sera[6,7]. Another study tested SARS-CoV-2-S pseudoviruses bearing either the Wuhan reference strain or the UK spike protein with BNT162b2 vaccine-elicited sera showed a slightly reduced but overall largely preserved neutralizing titers against the UK pseudovirus[8]. However, none of these studies was performed on clinical isolates harboring the full genomic mutations background of UK and SA strains. Thus, the question remained whether a replicating virus with the full set of S mutations, which may potentially interfere with antibody binding would be neutralized efficiently by convalescent COVID-19 or BNT162b2-immune sera, especially in the healthcare workers (HCW), a particularly exposed population to SARS-CoV-2 infection.

To answer this question, we performed a virus neutralization test (VNT), with a strict 100% inhibition criterion, on sera from HCW with either previous mild forms of COVID-19 or BNT162b2 immunization using three clinical isolates of SARS-CoV-2 variants: a D614G strain (D614G) which became the dominant form of the virus circulating globally in the second part of 2020[2], a UK strain (UK, lineage B.1.1.7) and a SA strain (SA, lineage B.1.351).

## MATERIALS AND METHODS

### Study population and serum specimen

Convalescent sera were recovered six months after symptom’s onset from symptomatic healthcare workers with a positive RT-PCR result. BNT162b2-vaccine elicited sera were recovered three weeks after the first injection and seven days after the booster immunization. This retrospective study was carried out in accordance with the Declaration of Helsinki without addition to standard of care procedures. Data collection were declared to the Sorbonne Université Data Protection Committee under number 2020-025 in accordance with French law. Written informed consent for participation in this study was obtained from all participants.

### SARS-CoV-2 IgG immunoassay

SARS-CoV-2 anti-nucleocapsid (N) IgG were determined using a commercially available immunoassay kit (Alinity SARS-CoV-2 IgG assay, Abbott Laboratories) according to the manufacturer’s instructions.

### SARS-CoV-2 strains

SARS-CoV-2 clinical isolates D614G, UK and SA (GenBank accession number MW322968, MW633280 and MW580244 respectively) were isolated from SARS-CoV-2 RT-PCR confirmed patients. Viral stocks were generated and titrated by the limiting dilution assay allowing calculation of tissue culture infective dose 50% (TCID50) after one passage of isolates on Vero cells.

### Virus neutralization test

The neutralizing activity of the various serum specimen was assessed with a whole virus replication assay as previously describe (*9*). Microscopy examination on day 4 to assess the cytopathic effect (CPE). Neutralization antibody (NAb) titers are expressed as the highest serum dilution displaying 100% inhibition of the CPE. A same known positive control serum was added to each experiment to assess the reproductivity.

### Statistical analysis

Difference in distribution of NAb titer between UK strain or SA strain with the D614G strain was performed with a two-tailed Mann-Whitney-U test in GraphPad Prism 6.0. A probability value of p<0.05 was considered statistically significant.

## RESULTS

We studied two sets of serum samples from HCW: a convalescent group of 15 participants with SARS-CoV-2 proven infection on March 2020 and a vaccinated group of 29 participants without history of clinical COVID-19 and which were negative for SARS-CoV-2 anti-nucleocapsid (N) IgG. The median [IQR] age was 50 [40 – 58] years and 40% (6/15) were male for the convalescent group. The median age was 55 [49 – 58] and 32% (9/29) were male for the vaccinated group. Convalescent sera were collected 6 months after the symptom’s onset (184 [182 – 189] days). BNT162b2 vaccine-elicited sera were collected 3 weeks after the first injection (except for four HCW) and 7 days after the second injection of BNT162b2-vaccine. All the serum samples were tested for their neutralizing activity against SARS-CoV-2 D614G, UK and SA clinical strains. Three weeks after the first injection of the BNT162b2 vaccine, 52% (13/25) of HCW harbored neutralizing activity with a NAb titer ≥ 1:5 against the D614G strain, 24% (6/25) were neutralizing against the UK strain and only two (8%) had detectable NAbs against the SA strain (Figure 1 A, Table S1). Seven days after the booster immunization, all but one HCW displayed neutralizing activity against the three SARS-CoV-2 clinical strains with a median neutralizing titer of 1:160 [80 – 160] against the D614G strain, 1:40 [40 –80] against the UK strain and 1:20 [20 – 40] against the SA strain. The median neutralizing titers against UK and SA strains were significantly reduced compared to median NAb titer against the D614G strain 7 days after the second injection of BNT162b2 vaccine (respectively, p < 0.0001 and p < 0.0001) (Figure 1 B, Table S1). Six months after the symptom’s onset, all the 15 HCW of the convalescent group harbored neutralizing activity against the D614G strain (median neutralizing titer of 1:20 [1:10 – 1:40]) and the UK strain (median neutralizing titer 1:20 [1:5 – 1:20]) without statistical difference between the respective NAb titers (p = 0.40). However, only 60% (9/15) serum samples of these HCW displayed a neutralizing activity against the SA strain with a median titer of 1:5 (Figure 1 C, Table S2).

**Fig. 1.**
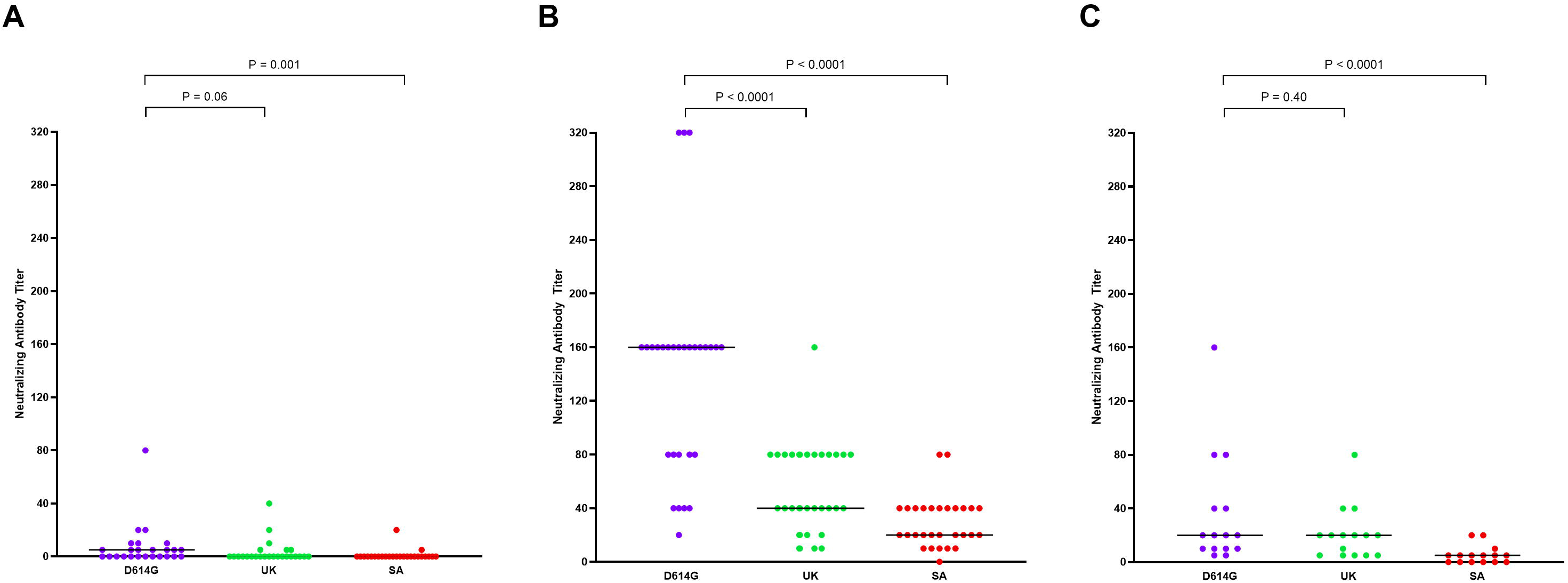
Neutralization antibody (NAb) titer against clinical strains of D614G, UK and SA SARS-CoV-2 variants of 29 BNT162b2-vaccine elicited sera and 15 convalescent sera recovered from healthcare workers (HCW). (A) NAb titer against the three clinical isolates of BNT162b2-vaccine elicited HCW sera recovered three weeks after first injection. (B) NAb titer against the three clinical isolates of BNT162b2-vaccine elicited HCW sera recovered seven days after second injection. (C) NAb titer against the three clinical isolates of convalescent COVID-19 HCW sera recovered 6 months after the symptom’s onset. NAb titer against D614G strain are in blue, NAb titer against UK strain are in red and NAb titer against SA strain are in green. Black horizontal lines indicate median values. Two-tailed P values were determined using the Mann-Whitney test and are reported on each panel.

## DISCUSSION

In this work we assessed the neutralizing activity of sera from 15 convalescent COVID-19 or 29 BNT162b2-vaccinated HCW against the two rapidly spreading SARS-CoV-2 variants of concern VOC202012/01 and 501Y.V2 and the globally circulating variant D614G using a VNT with whole replicating clinical strains. Based on a very strict criterion of 100% inhibition of CPE to determine NAb titer, we show that, three weeks after a single dose of BNT162b2, these NAb titers remain low or absent among HCW especially against the UK and SA variants and could questioned the extend of the dosing interval of BNT162b2 in some countries in order to vaccinate as many people as possible. However, we were not able to follow participants more than three weeks after the first injection because all of them received a second dose of BNT162b2 according to the French guidelines. Nevertheless, seven days after the booster immunization all but one vaccinated HCW develop NAbs against the three strains with a highest neutralizing activity against the strain closely related to the Wuhan ancestral strain, the D614G strain. Despite lower NAb titers against the UK and the SA strains most of the participants have displayed a neutralizing activity ≥ 1:10 which could be at least indicative of a potential protection against severe COVID-19 even with these variants. We also demonstrate a lack of serum neutralizing activity against SA strain in up to 40% of HCW recovered from mild form of COVID-19 six months after the symptom’s onset. This finding, and the recent report describing a severe case of reinfection by the SA variant four months after a first COVID-19 infection[9], highlights the need of vaccination even in people who had recovered from a previous COVID-19, especially during the increased circulation of the SARS-CoV-2 variants. Nevertheless, correlates of immunity to the SARS-CoV-2 are not well defined, only few studies have tried to assess these correlates in other human coronaviruses (HCoV) with experimental challenges on volunteers. They showed an association between serum NAb titer pre-exposure and viral excretion[10]. Further studies are required to determine the SARS-CoV-2 correlates of vaccine-induced protection based on NAb and T cell responses. A limitation of our work is that we were not able to assess potential cellular response differences against the three strains in the vaccinated or convalescent groups although it has been described generation of a robust CD4+ and CD8+ responses against the Wuhan ancestral strain[11]. The long-term evaluation regarding the lasting of NAb induced by vaccination is needed to assess the durability of protection against SARS-CoV-2 variants.

## CONCLUSION

In conclusion, in BNT162b2-vaccinated participants with two dose regimen, despite heterogeneity neutralizing capacity against the three SARS-CoV-2 variants, most of the sera harbored at least a NAb titer ≥ 1:10. Although immune protection correlates need to be defined, our findings suggests a certain humoral protection activity either on UK or SA variants after two doses of mRNA-vaccine. However, we show that six months after SARS-CoV-2 infection leading to mild forms of COVID-19, an important proportion of HCW displayed no neutralizing activity against SA strain. This result supports a strong recommendation for SARS-CoV-2 vaccination of previously infected subjects.

## Acknowledgments

We thank all the members of the Pitié-Salpêtrière Virology Department for their active collaboration and Vanessa Demontant, Elisabeth Trawinski, Melissa N’Debi and Guillaume Gricourt for helpful technical assistance in the full genome sequencing of the UK and SA strains.

---

**Table S1.** Serum neutralizing antibody (NAb) titer against three clinical strain of 29 vaccinated healthcare workers three weeks after the first dose of 30 μg BNT162b2 and seven days after the second dose. D614G: globally circulating D614G variant, UK: VOC202012/01 variant and SA: 501Y.V2 variant. N/A: non-applicable, four serums were not available for testing three weeks after the first injection of BNT162b2.

| Serum ID | 3 weeks after first BNT162b2 injection<br>(NAb titer) |  |  | 7 days after second BNT162b2 injection<br>(NAb titer) |  |  |
| --- | --- | --- | --- | --- | --- | --- |
|  | D614G strain | UK strain | ZA strain | D614G strain | UK strain | ZA strain |
| V1 | 10 | 0 | 0 | 160 | 40 | 20 |
| V2 | 5 | 0 | 0 | 160 | 40 | 40 |
| V3 | 5 | 0 | 0 | 160 | 40 | 20 |
| V4 | 20 | 40 | 5 | 160 | 80 | 40 |
| V5 | 0 | 0 | 0 | 80 | 40 | 20 |
| V6 | 0 | 0 | 0 | 40 | 10 | 10 |
| V7 | 0 | 0 | 0 | 160 | 40 | 10 |
| V8 | 0 | 0 | 0 | 40 | 20 | 10 |
| V9 | 0 | 5 | 0 | 40 | 20 | 10 |
| V10 | 0 | 0 | 0 | 160 | 80 | 20 |
| V11 | 0 | 0 | 0 | 160 | 40 | 40 |
| V12 | 0 | 0 | 0 | 160 | 80 | 40 |
| V13 | 5 | 0 | 0 | 160 | 40 | 20 |
| V14 | 5 | 0 | 0 | 80 | 40 | 20 |
| V15 | 5 | 5 | 0 | 160 | 80 | 40 |
| V16 | 5 | 0 | 0 | 160 | 40 | 40 |
| V17 | N/A | N/A | N/A | 160 | 80 | 80 |
| V18 | 5 | 5 | 0 | 80 | 20 | 40 |
| V19 | 0 | 0 | 0 | 80 | 80 | 20 |
| V20 | 0 | 0 | 0 | 160 | 80 | 40 |
| V21 | 10 | 20 | 0 | 320 | 160 | 20 |
| V22 | 0 | 0 | 0 | 160 | 80 | 40 |
| V23 | 80 | 10 | 20 | 320 | 80 | 80 |
| V24 | 10 | 0 | 0 | 160 | 80 | 20 |
| V25 | 20 | 0 | 0 | 160 | 80 | 20 |
| V26 | 0 | 0 | 0 | 20 | 10 | 0 |
| V27 | N/A | N/A | N/A | 80 | 40 | 40 |
| V28 | N/A | N/A | N/A | 40 | 10 | 10 |
| V29 | N/A | N/A | N/A | 320 | 80 | 40 |

**Table S2.** Serum neutralizing antibody (NAb) titer against three clinical strains of 15 COVID-19 recovered healthcare workers six months after symptom’s onset. D614G: globally circulating D614G variant, UK: VOC202012/01 variant and SA: 501Y.V2 variant.

| Serum ID | D614G strain | UK strain | ZA strain |
| --- | --- | --- | --- |
| I1 | 20 | 10 | 5 |
| I2 | 5 | 5 | 0 |
| I3 | 5 | 5 | 0 |
| I4 | 10 | 5 | 0 |
| I5 | 10 | 20 | 0 |
| I6 | 20 | 5 | 5 |
| I7 | 10 | 5 | 0 |
| I8 | 10 | 20 | 5 |
| I9 | 20 | 20 | 5 |
| I10 | 40 | 20 | 5 |
| I11 | 80 | 80 | 20 |
| I12 | 40 | 40 | 0 |
| I13 | 20 | 20 | 5 |
| I14 | 80 | 20 | 10 |
| I15 | 160 | 40 | 20 |

